# Prevalence Estimates of Putatively Pathogenic Leptin Variants in the gnomAD Database

**DOI:** 10.1101/2022.03.25.485774

**Authors:** Luisa Sophie Rajcsanyi, Yiran Zheng, Pamela Fischer-Posovszky, Martin Wabitsch, Johannes Hebebrand, Anke Hinney

**Affiliations:** Department of Child and Adolescent Psychiatry, Psychosomatics and Psychotherapy, University Hospital Essen, University of Duisburg-Essen, Essen, Germany; Center for Translational Neuro- and Behavioural Sciences, University Hospital Essen, Essen, Germany; Division of Pediatric Endocrinology and Diabetes, Department of Pediatrics and Adolescent Medicine University Medical Center Ulm, Ulm, Germany

**Author notes:** Corresponding author, (AH).

## Abstract

Homozygosity for pathogenic variants in the leptin gene leads to congenital leptin deficiency causing early-onset extreme obesity. This monogenic form of obesity has mainly been detected in patients from consanguineous families. Prevalence estimates for the general population using the Exome Aggregation Consortium (ExAC) database reported a low frequency of leptin mutations. One in approximately 15 million individuals will be homozygous for a deleterious leptin variant. With the present study, we aimed to extend these findings utilizing the augmented Genome Aggregation Database (gnomAD) v2.1.1 including more than 140,000 samples. In total, 68 non-synonymous and 7 loss-of-function (LoF) leptin variants were deposited in gnomAD. By predicting functional implications with the help of *in silico* tools, like SIFT, PolyPhen2 and MutationTaster2021, the prevalence of hetero- and homozygosity for putatively pathological variants (n = 32; pathogenic prediction by at least two tools) in the leptin gene were calculated. Across all populations, the estimated prevalence for heterozygosity for functionally relevant variants was approximately 1:2,100 and 1:17,860,000 for homozygosity. This prevalence deviated between the individual populations. Accordingly, people from South Asia were at greater risk to carry a possibly damaging leptin variant than individuals of other ancestries. Generally, this study emphasises the scarcity of deleterious leptin variants in the general population with varying prevalence for distinct study groups.

## Introduction

The leptin-melanocortin system modulates energy homeostasis and body weight regulation via the hypothalamic arcuate nucleus (ARC). The hormone leptin (LEP) is secreted by the adipose tissue into the bloodstream. In the ARC, leptin binds to the leptin receptor on pro-opiomelanocortin (POMC) and agouti-related peptide (AgRP) expressing neurons, stimulating POMC release and inhibiting AgRP expression. Subsequently, POMC is post-translationally processed into e.g. the α-melanocyte-stimulating hormone. Eventually, the signalling of melanocortin-4-receptor is stimulated and leads to decreased food intake due to satiety signals (1–5).

Homozygous mutations in the *LEP* gene cause congenital leptin deficiency disrupting the normal regulation of the body weight. Leptin levels in homozygous carriers of deleterious mutations are in most cases extremely low to undetectable (5, 6). Some deleterious mutations lead to a biologically inactive leptin. Leptin levels in these patients are seemingly normal for their BMI (6). A rapid weight gain eventually leads to extreme obesity. Hyperphagia, hypogonadism and impaired immune functions are concomitant symptoms (5, 7–9, 10). This form of monogenic obesity is infrequent, with a prevalence between one and 5% and predominantly affecting individuals with parental consanguinity (5, 7, 8, 11–14). In 1997, the first deleterious *LEP* mutation (p.Gly133Val*fs*\*15) was reported by Montague and colleagues (7). It was detected in the homozygous state in two cousins descending from a consanguineous family with the unaffected parents being heterozygous carriers of the variant. Due to this frameshift mutation, the LEP protein was truncated as 14 aberrant amino acids and a premature stop codon were introduced. This led to a rapid onset of obesity after normal birth weight (7). Subsequent treatment with recombinant leptin led to a substantial weight loss and a decrease in energy intake (11, 15). Further, besides frameshift mutations, pathogenic nonsense, and non-synonymous variants as well as deletions in *LEP* have been reported (5). The functional effects of these mutations are diverse. For instance, a deletion (p.Ile35del) that has been detected in one homozygous obese patient leads to a complete loss of the second exon of *LEP* and the removal of an isoleucine from the N-terminus of the protein (16). Interestingly, the non-synonymous variant p.Asp100Tyr was detected in an extremely obese boy from a consanguineous family. He showed high serum leptin levels and a pronounced history of infections. Functional analyses revealed normal leptin expression and secretion but a dysfunctional bio-inactive leptin that did not induce Stat3 phosphorylation (6, 14).

In 2017, Nunziata et al. (17) estimated the prevalence of putatively damaging mutations in the *LEP* gene using the Exome Aggregation Consortium (ExAC) database. Based on data from 60,706 samples, it was estimated that one in 15,000,000 individuals is potentially a homozygous carrier of a deleterious *LEP* mutation (determined by *in silico* tools), while approximately one in 2,000 individuals harbours a heterozygous leptin variant (17). Upon inclusion of functionally relevant *LEP* variants described in the literature, the authors estimated higher prevalence of hetero- and homozygosity of 1:1,050 and 1:4,400,000, respectively (17). To date, ExAC has been augmented into the Genome Aggregation Database (gnomAD) including more than 140,000 samples (version v2.1.1; 18). Therefore, we aimed to estimate the prevalence of putatively deleterious non-synonymous, frameshift and nonsense (loss-of-function: LoF) mutations in the *LEP* gene based on this extended dataset represented in gnomAD v2.1.1.

## Methods

### gnomAD

The gnomAD database (https://gnomad.broadinstitute.org/, accessed: Jan 24^th^, 2022), encompasses 15,708 full genome and 125,748 exome sequencing datasets from individuals of diverse populations (v2.1.1, GRCh37/hg19) comprising more than 200 million genetic variants. The sequencing data predominantly originates from case-control studies of diseases diagnosed in adulthood, such as cardiovascular diseases or psychiatric disorders. To ensure high quality data, all samples were subjected to a quality control, excluding samples with low sequencing quality, samples from second-degree relatives or higher, and data from patients with severe pediatric diseases. In total, six global and eight sub-continental populations are included, while populations from the Middle East, Central and Southeast Asia, Oceania and Africa are generally underrepresented. The mean coverage of the *LEP* gene was ~ 80x for exome and ~ 30x for genome data (18).

### Leptin Variants and their Predicted Functional Implications

In gnomAD, the *LEP* gene (canonical transcript ENST00000308868.4) was analysed and data pertaining to non-synonymous and LoF variants as well as the corresponding population-specific allele counts, and frequencies were extracted.

Consequences on the leptin protein by non-synonymous variants were predicted utilizing various *in silico* tools, namely Sorting Intolerant From Tolerant (SIFT, 19), Polymorphism Phenotyping v2 (PolyPhen2, 20), MutationTaster2021 (21), Functional Analysis through Hidden Markov Models – multiple kernel learning (FATHMM-MKL, 22) and Protein Variation Effect Analyzer (PROVEAN, 23). Predictions by SIFT, FATHMM-MKL and PROVEAN were obtained with the help of the Variant Effect Predictor (VEP, 24). For LoF, gnomAD presents predictions whether the respective LoF variant is a high- or low-confidence LoF based on results of either the LOFTEE tool or a manual curation, shown below the information of VEP on gnomAD’s variant page (18, 25).

SIFT classifies variants as either ‘tolerated’ or ‘deleterious’, while PolyPhen2 categorizes the mutations into ‘benign’, ‘possibly damaging’ and ‘probably damaging’. For PolyPhen2, the ‘HumVar’ classifier model was applied. MutationTaster2021 subjects each variant to several *in silico* tools itself and subsequently annotates each substitution as either ‘benign’ or ‘deleterious’. FATHMM-MKL and PROVEAN classify the variants into two categories: ‘neutral’ and ‘damaging’. Except MutationTaster2021, all these tools exclusively analyse non-synonymous variants. Thus, we annotated frameshift mutations with the LoF confidence predictions stated on the variant’s page.

Based on the preceding *in silico* analyses, the probability of hetero- and homozygous variants predicted to be pathogenic was calculated applying the Hardy-Weinberg equilibrium with the assumption of a perfect population (see Equation (1); p = allele frequency of allele A, q = allele frequency of allele a).

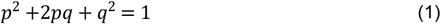

Hence, the prevalence of the heterozygous (*2qp*) and homozygous (including compound heterozygous; *q^2^*) variants were determined (see Equation (1)). To assess the prevalence of homozygous variants, the frequencies (*q^2^*) of the individual alleles were calculated and subsequently summed up. Subtraction of the prevalence of homozygosity from the prevalence of homozygous including the compound heterozygous variants revealed the corresponding frequencies for the compound heterozygotes.

When analysing the individual populations, substitutions were considered pathogenic if at least two of the applied *in silico* tools identified the variant as ‘damaging’ or ‘deleterious’ of if it was a high-confidence LoF variant.

## Results

In total, 75 non-synonymous and LoF variants in the *LEP* gene were deposited in gnomAD. Of these, 68 were non-synonymous (90.70%), five were frameshift (6.67%) and one each was an in-frame deletion (1.33%) or splice acceptor variant (1.33%). Across all populations, the non-synonymous variant rs17151919 (p.Val94Met) was the most frequent polymorphism with an overall allele frequency (AF) of 0.84% (see Table 1). A total of 105 homozygous and 2,167 heterozygous carriers of rs17151919 were observed (see S1 Table). Yet, *in silico* tools predicted a non-pathogenic potential (see S1 Table).

**Table 1:** Summary of non-synonymous and LoF variants in the *LEP* gene as deposited in gnomAD.

| Population | Sample size | Total number of variants* | Number of non-synonymous variants | Number of LoF variants | Most common variant (AF) |
| --- | --- | --- | --- | --- | --- |
| All populations | 141,456 | 75 | 68 | 7 | rs17151919<br>(0.84%) |
| African-American | 12,487 | 13 | 12 | 1 | rs17151919<br>(8.41%) |
| Ashkenazi Jewish | 5,185 | 1 | 1 | 0 | rs17151919<br>(0.26%) |
| East-Asian | 9,977 | 14 | 13 | 1 | rs148407750<br>(0.31%) |
| European, Finnish | 12,562 | 3 | 3 | 0 | rs751272426<br>(0.04%) |
| European, non-Finnish | 64,603 | 41 | 37 | 4 | rs17151919<br>(0.04%) |
| Latino/Admixed American | 17,720 | 14 | 14 | 0 | rs17151919<br>(0.45%) |
| **Others** | 3,614 | 8 | 8 | 0 | rs17151919<br>(0.38%) |
| **South Asian** | 15,308 | 16 | 14 | 2 | rs17151919<br>(0.03%) |

Table 1 summarizes the number of *LEP* variants (non-synonymous and LoF\*) deposited in gnomAD. The most common variants detected in various populations were predicted to be benign (see S1 Table). AF: allele frequency. LoF: loss-of-function.

Considering the populations individually, the aggregation of samples of multiple smaller populations denominated as ‘Other’ showed the highest occurrence of non-synonymous and LoF variants when correcting for the respective population size (0.0022; eight variants in total; see Table 1). Occurrence rates of these variants in populations from East and South Asian countries were lower with 0.0014 (total of 14 variants) and 0.0010 (total of 16 variants), respectively. Within the African-American population, non-synonymous and LoF variants showed a population size-corrected frequency of 0.00104 (total of 13 variants). Lower occurencies were detected in the Latino/Admixed population (0.0008; total of 14 variants), the European, non-Finnish (0.0006; total of 41 variants), the European, Finnish (0.00024; total of three variants) and the Ashkenazi Jewish population (0.0002; solely one variant; see Table 1). Generally, for all populations, the majority of variants was annotated as non-synonymous mutations (> 85%) and were rare (AF < 1%, see Table 1). Solely, the non-synonymous and putatively benign single nucleotide polymorphism (SNP) rs17151919 (see S1 Table) was frequent in African-Americans with an AF of 8.4%. Further, this SNP was the most commonly detected variant in European, non-Finnish individuals as well as in the African-American, Latino/Admixed American, Ashkenazi Jewish, South Asian and ‘other’ populations (see Table 1). Conversely, in Finnish samples, the variant rs751272426 (AF = 0.0047%) was the most common, while rs148407750 (AF = 0.311%) was the most abundant SNP in people from East Asia.

To assess functional implications of the *LEP* variants in gnomAD, we analysed the variants with various *in silico* tools (see S1 Table). Accordingly, SIFT assigned nine variants as ‘deleterious’, while PolyPhen2 predicted 13 variants to be ‘possibly damaging’ and 16 to be ‘probably damaging’. Ten variants were assigned as ‘deleterious’ by MutationTaster2021. ‘Damaging’ classifications for 24 and 20 variants were obtained by FATHMM-MKL and PROVEAN, respectively (see Fig. 1 and S1 Table). Additionally, five of six LoF variants were indicated to be high-confidence LoF variants (see S1 Table). Twenty-two variants across all populations were predicted to be benign (see Table 2 and S1 Table). Fifty-three variants were indicated to be pathogenic by at least one tool, while 32 and 19 revealed a deleterious effect in at least two and three tools, respectively. Collectively, one in approximately 53 individuals will be a carrier of a non-synonymous or LoF variant located in *LEP* regardless of the pathogenicity (see Table 2). The prevalence for a homozygous and compound heterozygous variant is ~ 1:14,100 and ~ 1:50,000, respectively. Lower prevalence were detected for variants predicted to be benign (see Table 2 and S1 Table). Generally, for indicated deleterious variants across all populations regardless of the number of tools predicting pathogenicity, the prevalence of compound heterozygosity is higher than the prevalence of homozygous variants (see Table 2). Further, when applying various pathogenicity definitions based on the number of *in* silico tools predicting a damaging effect, it is evident that the more stringent this definition, the lower the prevalence (see Table 2). Consequently, we decided to classify variants as pathogenic if at least two *in silico* tools indicated a deleterious impact (definition applied for subsequent analyses). A total of 67 individuals throughout all populations carried at least one of these variants heterozygous, while no homozygous carriers were detected. Hence, the estimated the estimated prevalence of heterozygosity for a putatively harmful *LEP* mutation was approximately 1:2,100, while the prevalence for a homozygous variant was ~1:17,860,000 for individuals of all populations (see Tables 2 and 3).

**Figure 1:**
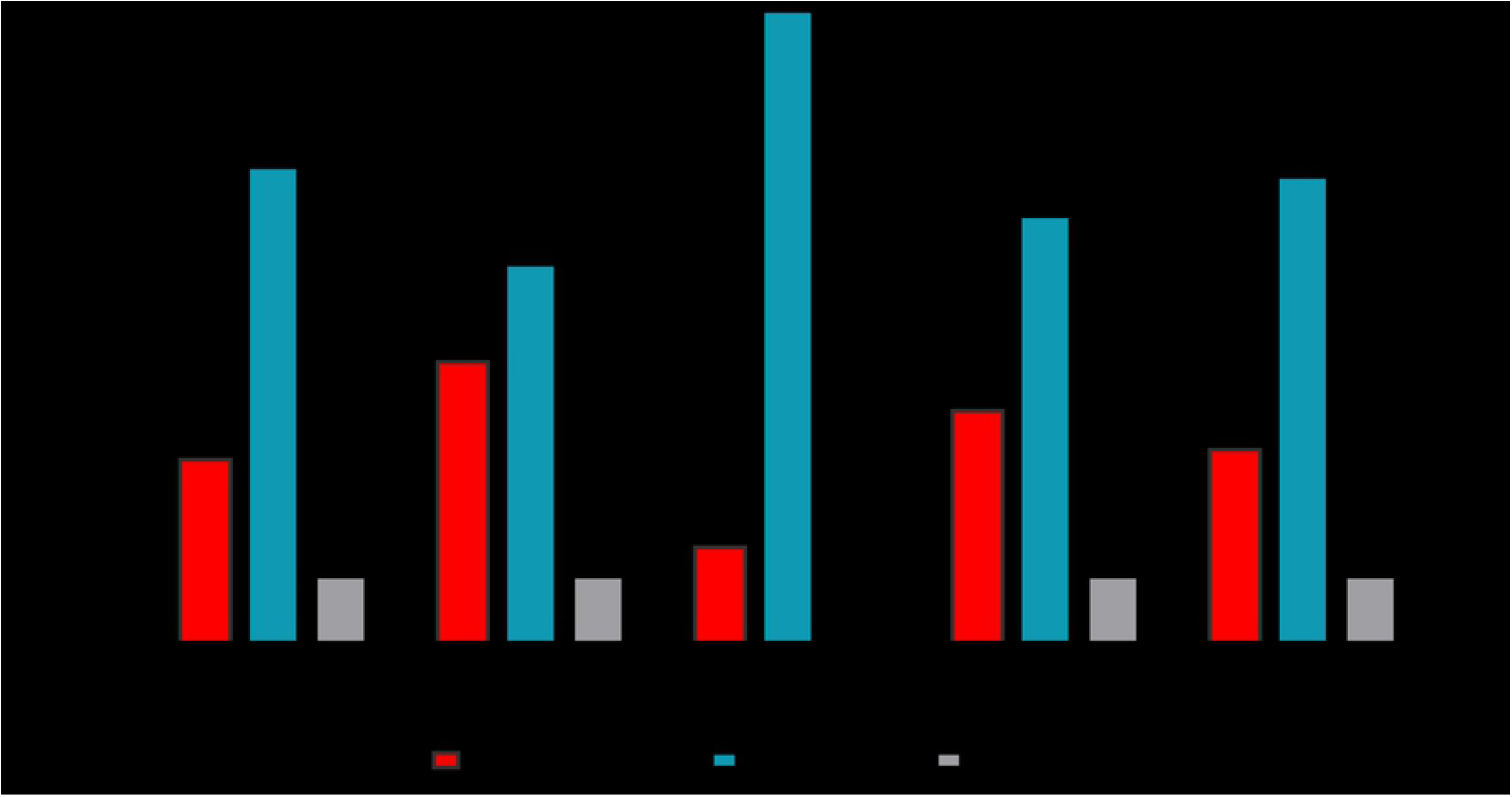
Predictions of the applied *in silico* tools. All 75 non-synonymous and LoF variants located in *LEP* were analysed with SIFT, PolyPhen2, MutationTaster2021, FATHMM-MKL and PROVEAN. Unless MutationTaster2021, all tools were unable to predict implications of the seven LoF variants (grey).

**Table 2:** Estimated prevalence of hetero- and homozygous as well as compound heterozygous variants across all populations.

| Number of tools predicting deleterious effect | Number of deleterious variants* | Number of carriers of deleterious variants |  | Estimated prevalence for heterozygous mutations | Estimated prevalence for homozygous and compound heterozygous mutations | Estimated prevalence of homozygous mutations | Estimated prevalence of compound heterozygous mutations |
| --- | --- | --- | --- | --- | --- | --- | --- |
|  |  | Heterozygous | Homozygous |  |  |  |  |
| 0 | 22 <sup>a</sup> | 2,254<sup>a</sup> | 105 <sup>a</sup> | 1 : 58 | 1 : 13,200 | 1 : 14,200 | 1 : 193,000 |
| ≥0 | 75 <sup>a</sup> | 2,486 <sup>a</sup> | 105 <sup>a</sup> | 1 : 53 | 1 : 11,000 | 1; 14,100 | 1 : 50,000 |
| ≥1 | 53 | 232 | 0 | 1 : 610 | 1 : 1,490,000 | 1 : 7,000,000 | 1 : 1,900,000 |
| ≥2 | 32 | 67 | 0 | 1 : 2,100 | 1 : 17,860,000 | 1 : 326,740,000 | 1 : 18,850,000 |
| ≥3 | 19 | 32 | 0 | 1 : 4,400 | 1 : 78,160,000 | 1 : 741,000,000 | 1 : 87,330,000 |

Here, the rounded estimated prevalence for variants in all populations are presented. Variants were considered deleterious if the stated number of *in silico* tools revealed a pathogenic prediction (\*). Due to varying allele frequencies across the individual populations, the Hardy-Weinberg-Equilibrium is not fulfilled when investigating exclusively the benign variants or all variants (<sup>a</sup>).

**Table 3:** Estimated prevalence for the populations in gnomAD.

| Population | Sample size | Number of putatively deleterious mutations* | Estimated prevalence for heterozygous mutations | Estimated prevalence for homozygous/compound heterozygous mutations |
| --- | --- | --- | --- | --- |
| All populations | 141,456 | 32 | 1 : 2,100 | 1 : 17,860,000 |
| All populations (females) | 64,754 | 19 | 1 : 2,200 | 1 : 18,640,000 |
| All populations (males) | 76,702 | 22 | 1 : 2,100 | 1 : 17,190,000 |
| African-American | 12,487 | 4 | 1 : 1,800 | 1 : 12,730,000 |
| Ashkenazi Jewish | 5,185 | 0 | NA | NA |
| East-Asian | 9,977 | 6 | 1 : 770 | 1 : 2,360,000 |
| European, Finnish | 12,562 | 0 | NA | NA |
| European, non-Finnish | 64,603 | 16 | 1 : 2,700 | 1 : 28,980,000 |
| Latino/Admixed American | 17,720 | 5 | 1 : 4,400 | 1 : 78,500,000 |
| Others | 3,614 | 5 | 1 : 900 | 1 : 3,270,000 |
| South Asian | 15,308 | 3 | 1 : 1,300 | 1 : 6,510,000 |

Estimated rounded prevalence for deleterious *LEP* variants are shown. Variants were considered deleterious if at least two *in silico* tools revealed a pathogenic prediction (\*).

gnomAD further provides sex-specific allele counts for each variant. Thus, we replicated the probability estimations of possibly pathogenic variants (as defined above) for both sexes separately. This revealed that about one in 2,200 women carries a heterozygous and possibly harmful *LEP* variant. In males, the prevalence of a heterozygous variant was marginally higher with ~1:2,100. The chance to harbour a homozygous/compound heterozygous, pathogenic variant in females was estimated to be ~1:18,640,000. For males, this prevalence was again higher at ~1:17,190,000.

Next, we determined the likelihood of a putatively deleterious *LEP* variant in the distinct populations. As none of the variants detected in the Finnish and Ashkenazi Jewish population was predicted to have a pathogenic effect, we were unable to calculate the correlated prevalence (see Table 3). Generally, pronounced variations between the global populations were observed (see Table 3). East Asians were determined to be at highest risk to harbour a putatively pathogenic *LEP* variant either hetero- or homozygous. The lowest risk for both hetero- and homozygous variants were found in the Latino/Admixed population (see Table 3).

## Discussion

Homozygous deleterious mutations in the leptin gene lead to deficiency of biologically active leptin and cause severe early onset obesity (5, 7, 8, 11, 14). Through the implementation of reference databases, such as ExAC and gnomAD, assessments of prevalence of potentially harmful variants in the general population have become feasible. Yet solely one study has explored the prevalence of *LEP* variants in the general population using these reference datasets (17). As of today, the gnomAD database is the largest publicly available databases containing data of genetic variants (25). More than 125,000 exome and 15,000 whole genome sequence datasets are contained in gnomAD v2.1.1 (18). Based on these datasets, it had been estimated that each individual carries approximately 200 coding variants with allele frequencies less than 0.1%. Despite the large sample size, gnomAD will lack on average 27 ± 13 novel coding mutations per exome based on the current number of samples included (25). The data contained in gnomAD has been subjected to a stringent quality control excluding data of participants with known severe pediatric diseases or related individuals (18, 25). Notably, due to this removal of samples with known pediatric diseases, potentially relevant and pathogenic variants with regard to early manifested obesity may have been omitted. Additionally, variation data regarding global cohorts are deposited in gnomAD. Still, European, non-Finnish samples are overrepresented, while samples from the Middle East, Central Asia and Africa are generally underrepresented. Since congenital leptin deficiency caused by mutations in leptin are more prevalent in patients from Pakistan and the Middle East (5, 7, 11, 16), there is a lack of data pertaining to deleterious leptin mutations in the general Middle Eastern population. It can be assumed that higher incidence of putatively harmful variants might be observed in these populations. Additionally, no individual-level phenotype data is available. Thus, it is unclear whether the datasets might be skewed for overweight or obese individuals, which is feasible considering the globally increasing prevalence of both (26).

We are aware that *in silico* tools are no substitute for functional *in vitro* analyses. This is particularly evident for the deletion p.Ile35del, as neither gnomAD, nor most *in silico* tools do provide predictions of functional implications. Yet, it is known that this deletion causes the loss of exon 2 of the *LEP* gene and thus a congenital leptin deficiency with resultant obesity (5, 16). Additionally, the performance of the individual tools varies considerably. A previous study has demonstrated that SIFT and PROVEAN yield the most accurate prediction of pathogenicity. MutationTaster2011 and FATHMM had comparatively low accuracy and specificity (27). Further, the more stringent the criteria of pathogenicity are defined, the lower the obtained prevalence (see Table 2). Accordingly, we classified variants as potentially harmful if at least two tools indicated a damaging effect. Generally, these tools do help to gain preliminary indications of putatively deleterious variants.

Across all populations, we detected that approximately one in 2,100 carries a potentially deleterious (when at least two *in silico* tools indicated a pathogenicity) heterozygous variant in *LEP*. In addition, the prevalence of a homozygous variant was about 1:17,860,000 among all populations. Despite the larger sample size, a resultant greater number of variants in gnomAD and a stricter definition of pathogenicity, our results resemble the estimated prevalence based on the ExAC database reported by Nunziata and colleagues (17). Thus, the here obtained prevalence of a harmful *LEP* variant in the general population confirms the low incidence of leptin variants reported in the previous study. Higher prevalence rates of hetero- and homozygosity are expected upon inclusion of functionally relevant variants reported in the literature (17). Generally, heterozygous variants were previously detected in unaffected individuals (5, 7). Heterozygous carries generally show lower BMI z-scores and lower body fat percentage than homozygous individuals (28). Yet, it is feasible that an additive effect of heterozygous variants may still have an impact on the carrier’s body weight. Throughout all populations, we report that compound heterozygous variants are less prevalent than single heterozygous variants but more frequent than homozygosity. In addition, we observed variations in prevalence rates between populations. For example, individuals from East Asia showed a higher prevalence of both heterozygous and homozygous mutations in *LEP* than other populations. Strong disparities were also evident at the SNP level. For example, the SNP rs17151919 was generally infrequently detected. In African-Americans, however, it was found frequently with an AF of well above 5%. One study has shown that the association of this SNP with lower leptin levels was specific for the African ancestry (29). Further, the functionally benign SNP rs17151919 was associated with a higher BMI in African children, but not in adults (29). Certain variants deposited in gnomAD and predicted to be pathogenic were previously associated with congenital leptin deficiency. The non-synonymous mutations p.Asp100Asn (rs724159998, 30), p.Asn103Lys (rs28954113, 6, 31, 32) and the frameshift mutation p.Gly133Val*fs*Ter15 (rs1307773933, 7) were detected in extremely obese children being homozygous carriers (5). Another study showed that BMI z-scores of carriers of homozygous *LEP* variants are generally higher than those of the heterozygous or wildtype individuals carrying the same variant (28).

## Conclusion

The gnomAD database is the largest publicly available reference dataset including various global study groups. By utilizing these datasets, we estimated the prevalence of putatively damaging leptin variants. We identified 19 possibly damaging mutations in 32 heterozygous and no homozygous carriers. The prevalence of a heterozygous variant was roughly 1:2,100, while the probability for homozygosity was 1:17,860,000 across all populations. Investigating each study group separately, this prevalence varied significantly, with East Asians being at greater risk of harbouring a hetero- or homozygous mutation with a harmful consequence. Yet, higher prevalence of functionally relevant variants could be obtained when reported case studies are included. In general, mutations in the *LEP* gene, which frequently result in congenital leptin deficiency, are extremely rare in the general population. Continued analysis of leptin mutations along phenotypic and clinical data may improve our understanding of monogenic obesity.

## Supporting Information Captions

**S1 Table: Summary of the results of the *in silico* analyses of *LEP* variants deposited in gnomAD.** Supplementary Table S1 represents the *in silico* predictions for each variant present in all populations by various tools, namely SIFT (19), PROVEAN (23), PolyPhen2 (20), MutationTaster2021 (21) and FATHMM-MKL (22). The tools are ordered by their reported accuracy (left: highest accuracy; right: lowest accuracy; 27).

